# Engineering the Structural Organization of Tryptophan in Crystalline Materials for Tunable Functionality

**DOI:** 10.64898/2026.07.26.740826

**Authors:** Oliver Ton, Sidhanth Duvvuri, Carol Korzeniewski, Raheleh Ravanfar

## Abstract

Tryptophan is a biologically important redox-active amino acid whose functions in proteins, including long-range electron transfer, protection against oxidative damage, and environmental sensing, are governed not only by its chemical identity but also by its precise structural organization. Inspired by this biological principle, we investigated whether controlling the organization of tryptophan within crystalline materials could provide a strategy for modulating its physicochemical properties and molecular accessibility. Using identical molecular components but distinct assembly pathways, tryptophan was organized either as a confined guest within a preformed Zn–imidazolate framework, yielding a star-shaped crystalline architecture, or as an integral coordination component during framework growth, producing a distinct layered Zn– tryptophan crystalline framework. Although assembled from the same building blocks, these two organization modes generated fundamentally different crystal structures, morphologies, and mechanisms of biomolecule incorporation. In both architectures, incorporation of tryptophan into the crystalline environment preserved its intrinsic fluorescence while producing robust fluorescence under multiple excitation wavelengths, highlighting the strong influence of molecular organization on its optical response. The structural modes also exhibited distinct encapsulation efficiencies and pH-dependent molecular accessibility, while secondary calcium–alginate fixation provided an additional level of control over guest retention without disrupting the underlying crystalline architecture. These results demonstrate that engineering the structural organization of tryptophan provides a versatile strategy for tuning the optical behavior, molecular accessibility, and functional integration of a biologically important redox-active amino acid in crystalline materials, establishing a foundation for future biomimetic redox architectures, responsive sensing platforms, and controlled molecular delivery.

## INTRODUCTION

Tryptophan (Trp) is among the most distinctive of the twenty canonical amino acids. Although it is the largest amino acid by molecular weight, it is also one of the least abundant, both as a constituent of proteins and as a free intracellular metabolite.^1–2^ This rarity is not a consequence of limited biological importance but rather reflects the selective placement of Trp residues within proteins, where they play key structural and functional roles.^3^ Evolutionary analyses have consistently identified Trp as one of the most highly conserved amino acids, indicating strong selective pressure to preserve its position within protein structures.^4^ Likewise, mutations involving Trp residues are disproportionately associated with human disease compared to most other amino acids, emphasizing their essential contributions to protein function.^5^ Consequently, Trp residues are rarely distributed randomly within proteins; instead, they are strategically positioned to fulfill specific structural and functional roles, including stabilizing protein structure, mediating intermolecular interactions, facilitating long-range electron transfer, protecting proteins against oxidative damage, and serving as intrinsically fluorescent reporters of the local molecular environment.^3, 6–11^ These unique physicochemical properties have stimulated growing interest in exploiting Trp as a functional molecular building block in synthetic materials for applications ranging from biosensing^12^ and drug delivery,^13–14^ to bioelectronics.^15–16^ However, despite increasing efforts to harness the intrinsic properties of Trp, comparatively little attention has been devoted to engineering its structural organization, even though its biological functions are fundamentally governed by its spatial arrangement within proteins. This dependence on structural organization is particularly evident in the intrinsic fluorescence of Trp, whose emission wavelength, intensity, and excited-state relaxation are exceptionally sensitive to the surrounding molecular environment, including polarity, hydrogen bonding, conformational restriction, and interactions with neighboring aromatic residues.^17–19^ More broadly, these observations highlight that the physicochemical properties of Trp emerge not solely from its molecular structure but also from its local structural environment, suggesting that controlling its organization within synthetic materials may provide a powerful strategy for modulating its function. Engineering the structural organization of Trp within synthetic materials requires a platform capable of precisely controlling its local coordination environment, spatial arrangement, and molecular accessibility. Metal– organic frameworks (MOFs) provide an attractive platform for this purpose owing to their well-defined crystalline architectures, tunable porosity, structural diversity, and synthetic versatility.^20–25^ Their ability to organize molecular building blocks within highly ordered frameworks has enabled the incorporation of a wide range of biomolecules while preserving their structural integrity and biological functionality for applications including drug delivery, stabilization, sensing, and catalysis.^26–32^

Here, we demonstrate that the structural organization of Trp within crystalline materials can be deliberately engineered through synthetic pathway selection. Using identical molecular building blocks under distinct assembly routes, Trp was organized either as a confined guest within a preformed Zn–imidazolate framework or incorporated directly during framework growth to generate a Zn–tryptophan crystalline framework in which the amino acid becomes an integral structural component (Figure 1, A-B). We find that these alternative organization modes produce fundamentally different crystal structures, morphologies, mechanisms of incorporation, encapsulation efficiencies, fluorescence behavior, and pH-dependent release. This organization-centered design was inspired by the way nature exploits the spatial arrangement of tryptophan residues to regulate biological function. In proteins, Trp residues participate in long-range Tyr/Trp hole-hopping pathways that protect against oxidative damage, where efficient electron transfer depends not only on the intrinsic redox properties of Trp but also on its precise spatial organization, local molecular environment, and solvent accessibility (Figure 1C).^6–8, 11, 33–34^ By establishing two complementary strategies for organizing tryptophan within crystalline materials, this work demonstrates that structural organization itself serves as an effective design parameter for modulating the physicochemical behavior of a biologically important redox-active amino acid, while providing a foundation for future investigations of biomimetic redox systems, responsive optical materials, and controlled molecular delivery.

**Figure 1.**
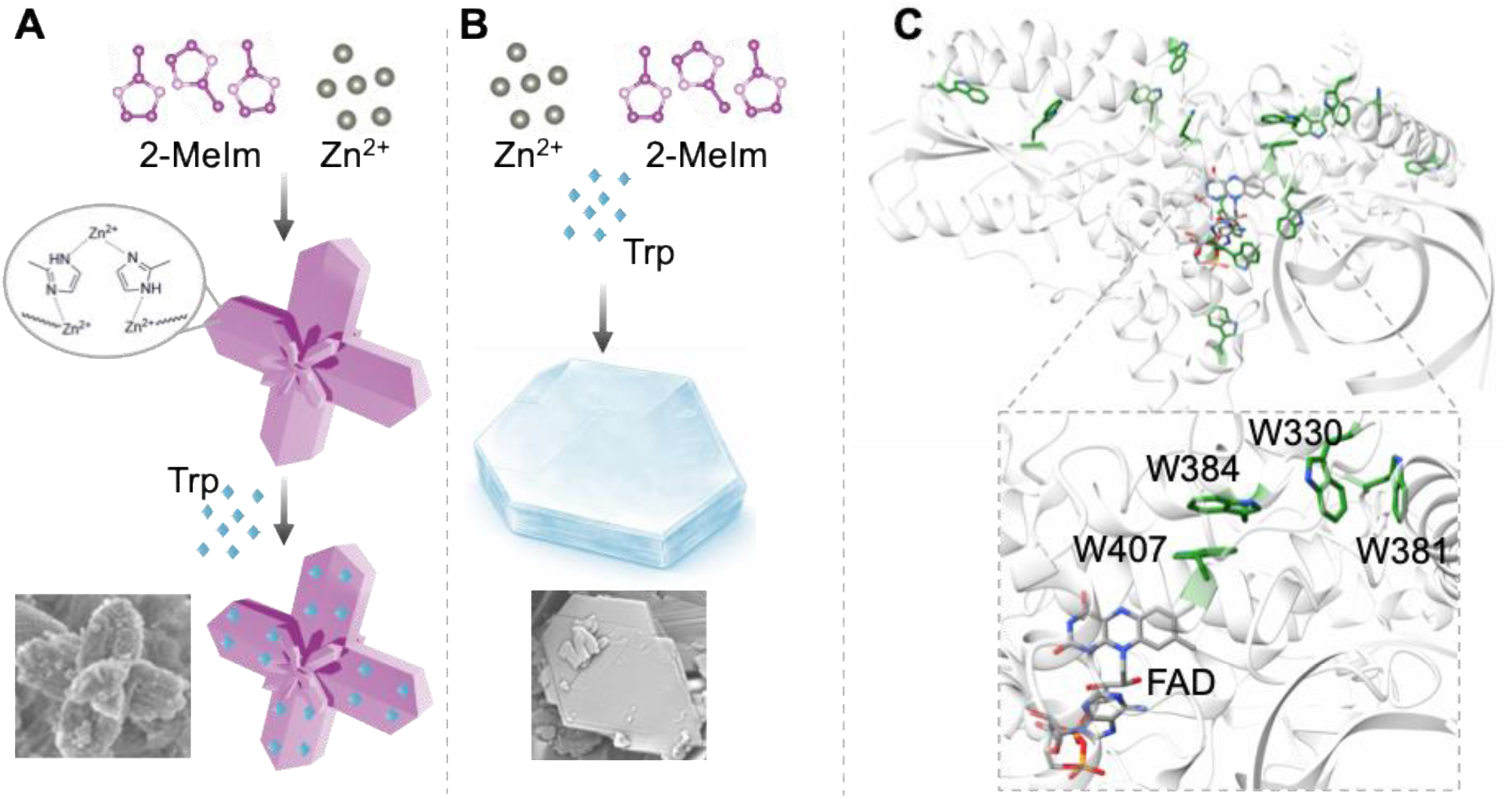
A) The schematic for post-synthetic incubation of Trp with preformed ZIF-8 OctaStars. B) The schematic for simultaneous combination of Zn^2+^, 2-MeIm, and Trp, generating a structurally distinct layered crystalline material. Representative SEM images highlight the pronounced morphological differences. C) Biological inspiration underlying the synthetic design.

## RESULTS AND DISCUSSION

Synthetic design enables direct comparison between guest confinement and coordination-driven integration. In the present study, two complementary synthetic routes were intentionally designed to isolate the role of incorporation pathway while maintaining identical molecular components throughout the system. Both materials were prepared using Zn^2+^, 2-methylimidazole (2-MeIm), and Trp, differing only in whether Trp was introduced before or after framework formation. The Trp’s indole moiety possesses multiple coordination sites capable of interacting with metal ions, allowing Trp to function either as a molecular guest or as a structural ligand during crystal growth. Such dual behavior provides a unique opportunity to investigate how different organization modes influence the resulting crystalline architecture and the physicochemical properties. In Route A, Zn–imidazolate particles were first synthesized independently to form faceted OctaStar architectures, after which Trp was incorporated post-synthetically through incubation to form OctaStar@Trp. In this pathway, Trp encounters an already-formed crystalline framework and therefore behaves primarily as a confined guest species interacting with the existing porous or interfacial environment (Figure 1A). In Route B, Zn^2+^, 2-MeIm, and Trp were combined simultaneously in a single one-pot synthesis, allowing Trp to compete with the canonical linker for the metal node and participate directly in coordination chemistry, nucleation, and crystal growth to form Trp-MOF. Under these conditions, Trp is no longer restricted to guest-like behavior and instead becomes involved in the structural organization of the resulting material (Figure 1B).

The structure of *Drosophila melanogaster* (6-4) photolyase (PDB Code: 3CVU), illustrating the spatial organization of Trp residues that mediate long-range electron transfer to photoexcited FAD. The inset demonstrates the Trp tetrad responsible for the electron-transfer pathway. FAD is shown in gray (colored by elements), and Trp residues are highlighted in green.

Single-crystal X-ray diffraction analysis revealed formation of a layered Zn–Trp coordination architecture of Trp-MOF, in which Zn centers are bridged through tryptophan-derived coordination motifs rather than classical imidazolate linkages (Figure S1). Trp-MOF crystalized in the monoclinic system with the space group of P2_1_ (Table S1). Each Zn center is five-coordinate, binding three oxygen atoms and two nitrogen atoms supplied by three connected Trp ligands. The resulting coordination environment is best described as distorted trigonal-bipyramidal. Two Trp ligands coordinate to Zn through both the amino nitrogen and a carboxylate oxygen atom, whereas a third Trp ligand coordinates through a carboxylate oxygen atom. Because the Trp ligands also connect neighboring Zn centers, this coordination motif propagates to generate an extended two-dimensional Zn-Trp coordination network (Figure S1, Table S1). This crystallographic analysis of the Route B material provides important mechanistic insight into the origin of the distinct diffraction behavior observed in comparison to both ZIF-8 and the Route A guest-confinement systems (Figure 2, A-B). The resolved structure of Route B consists predominantly of Zn–O/N coordination environments associated with Trp ligands, producing an extended layered organization stabilized through both coordination bonding and supramolecular interactions between neighboring aromatic indole groups. The indole nitrogen does not participate in Zn coordination because its lone pair is delocalized within the aromatic π system, making it a poor Lewis base compared with the amino and carboxylate groups. Importantly, although 2MeIm was required during synthesis to generate the crystalline layered MOF in Route B, it was not observed as a coordinated ligand within the final resolved crystal structure (Figure S1). This observation strongly suggests that 2MeIm acts as a transient assembly mediator, likely influencing pH, Zn coordination equilibria, nucleation, or crystal-growth kinetics, rather than serving as a permanent linker in the final lattice. This interpretation is consistent with the broader concept of modulated MOF/coordination-solid synthesis, where additives or competing ligands can control crystallinity, particle size, morphology, phase selection, and kinetic assembly pathways without necessarily defining the final framework connectivity.^35–37^

**Figure 2.**
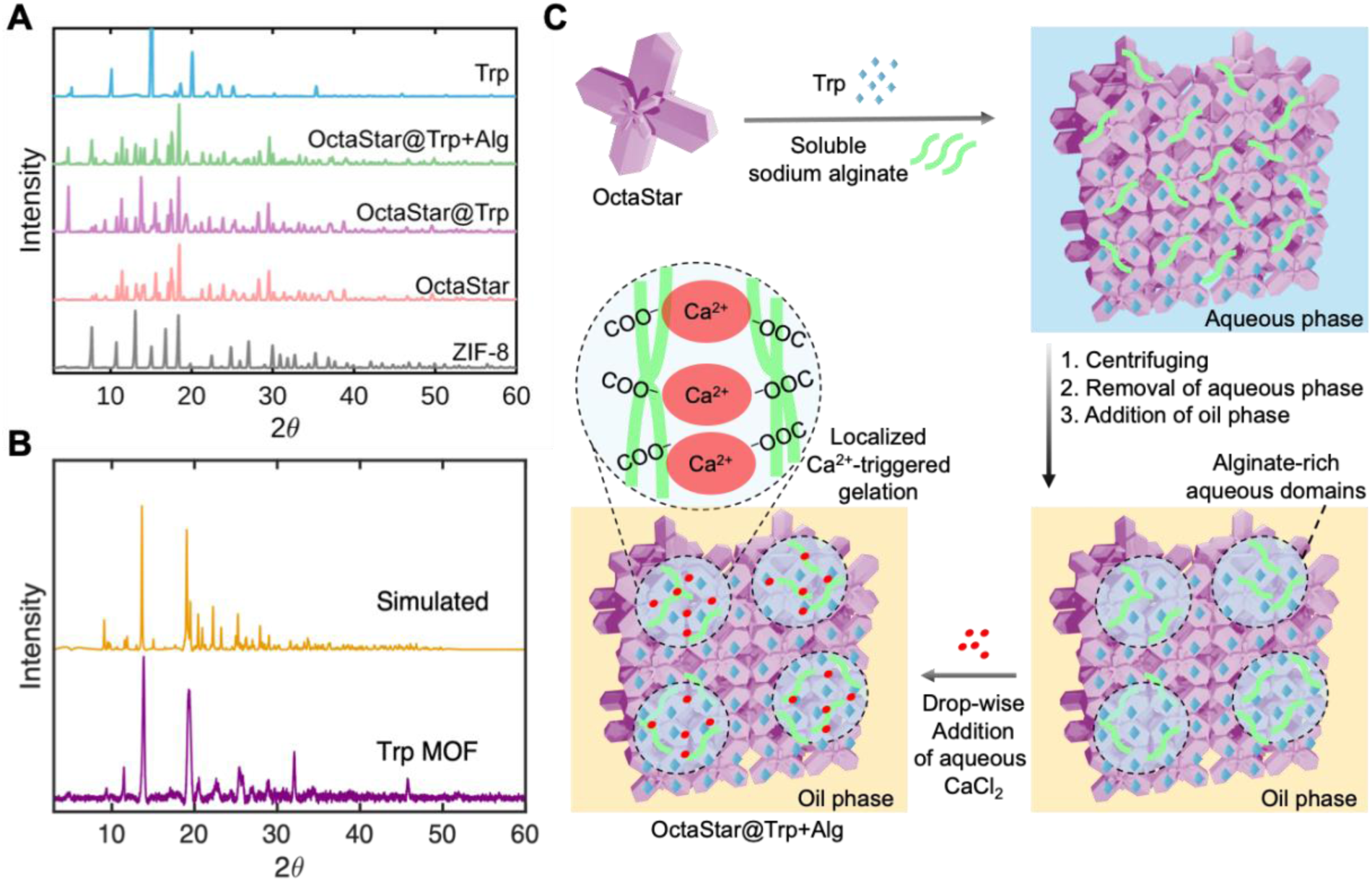
Structural divergence arising from pathway-dependent incorporation of tryptophan into Zn-containing MOFs. A) PXRD patterns of Route A materials, including ZIF-8, OctaStar, OctaStar@Trp, and OctaStar@Trp/Alg, together with free Trp. B) PXRD patterns of the Route B material (Trp-MOF, purple) and the simulated diffraction pattern generated from the single-crystal X-ray structure (yellow). The agreement between experimental and simulated patterns confirms the formation of a distinct crystalline phase that differs substantially from the Zn–imidazolate frameworks observed in Route A. C) Schematic illustration of OctaStar@Trp+Alg formation through localized Ca^2+^-triggered gelation of Trp-containing alginate domains within and around the OctaStar voids.

This structural result explains the pronounced difference between the PXRD pattern of the Route B Trp-MOF (Figure 2B) and that of ZIF-8 (Figure 2A, gray spectrum). Classical ZIF-8 is built from tetrahedral Zn centers bridged by 2-MeIm linkers to form a sodalite-type zeolitic framework, giving rise to a characteristic diffraction pattern.^38–39^ In contrast, the Route B Trp-MOF exhibits a different set of reflections and intensity distribution, confirming that its crystallographic organization does not correspond to conventional ZIF-8 or to simple Trp adsorption on a ZIF-type phase (Figure 2B). Importantly, the experimental PXRD pattern of the Route B MOF agrees well with the simulated pattern generated from the single-crystal structure (Figures 2B & S1), supporting the assignment of the bulk material to the layered Zn–Trp coordination phase in Trp-MOF (Figure 2B). This agreement is critical because it links the single-crystal model to the powdered sample and argues against the material being merely a physical mixture of Trp and a Zn–imidazolate phase.

In contrast to the coordination-driven assembly observed in Trp-MOF in Route B, the PXRD data obtained for OctaStar@Trp in Route A reveal a fundamentally different mode of tryptophan incorporation (Figure 2A). Notably, the diffraction pattern of OctaStar differs substantially from that of conventional ZIF-8, indicating that the material does not correspond to the canonical sodalite topology typically associated with ZIF-8.

Although both materials are synthesized from Zn^2+^ and 2-MeIm, their synthetic conditions differ significantly, particularly in the ligand-to-metal ratio. OctaStar was prepared using a Zn:2-MeIm ratio of approximately 1:16, whereas conventional ZIF-8 was synthesized using a substantially larger excess of ligand (Zn:2-MeIm ∼ 1:60). Previous studies have demonstrated that the metal-to-ligand ratio strongly influences nucleation kinetics, crystal growth rates, morphology, and phase evolution in Zn-imidazolate systems, often leading to the formation of distinct Zn-imidazolate architectures under otherwise similar reaction conditions.^40–41^ The lower ligand excess employed during OctaStar synthesis may therefore alter the balance between nucleation and crystal growth, promoting formation of the unique star-like phase observed via Route A (Figure 2A). Interestingly, several characteristics of OctaStar appear indicative of layered Zn-imidazolate materials such as ZIF-L.^42–44^ Unlike the 3D sodalite framework of ZIF-8, ZIF-L consists of layered Zn-imidazolate sheets stabilized by hydrogen-bonded neutral imidazole species and exhibits anisotropic crystal growth behavior.^42^ While the diffraction pattern of OctaStar does not fully match reported ZIF-L references and therefore cannot be definitively assigned as a conventional ZIF-L phase, its distinct diffraction profile and anisotropic morphology suggest that it may represent a related Zn-imidazolate architecture formed under kinetically controlled growth conditions. Regardless of its precise structural assignment, the OctaStar phase clearly differs from classical ZIF-8 and provides a unique host lattice for subsequent biomolecule incorporation. Importantly, post-synthetic incubation of OctaStar with Trp does not induce substantial reconstruction of the crystalline framework. Instead, the dominant diffraction features of the parent OctaStar phase remain largely preserved following Trp incorporation in OctaStar@Trp (Figure 2A).

Post-synthetic incorporation provides a straightforward route for introducing Trp into the preformed OctaStar host, but guest confinement alone may not ensure long-term retention of the amino acid within the material. We therefore investigated a secondary fixation strategy based on calcium alginate gelation to further stabilize the guest and restrict Trp release from the OctaStar host (Figure 2C). To achieve this hierarchical organization, preformed OctaStars were exposed to Trp in the presence of soluble sodium alginate, allowing guest incorporation and association of the polysaccharide before ionic cross-linking (Figure 2C). Upon centrifugation, the particles were recovered from the aqueous phase and dispersed in an oil phase, followed by gradual introduction of Ca^2+^ to induce alginate gelation and ultimate formation of OctaStar@Trp+Alg (Figure 2C). In this design, the oil phase does not directly participate in alginate cross-linking; instead, it serves as a hydrophobic continuous phase that restricts the spatial distribution of the aqueous alginate-containing domains and thereby promotes localized gel formation within or around the OctaStar voids. This sequence minimizes the presence of freely dissolved alginate in a continuous aqueous phase, thereby reducing the likelihood of independent bulk alginate gel or bead formation. Upon gradual addition of aqueous CaCl_2_, the strong partitioning preference of the hydrated Ca^2+^ ions for residual water-rich domains is expected to favor their transport toward localized hydrophilic regions containing alginate and Trp within or around the OctaStar voids (Figure 2C). Alginate gelation occurs through ionic association of Ca^2+^ with carboxylate-rich regions of neighboring polysaccharide chains, particularly guluronate-rich sequences, generating intermolecular junction zones commonly described by the egg-box model.^45–46^ This gelation process provides a mild, and biomolecule-compatible route for immobilization without requiring covalent modification of the confined guest. Oil-phase or emulsion-assisted approaches are widely used to spatially confine alginate gelation and produce alginate-based particulate or compartmentalized materials.^47^ Following alginate treatment, the principal reflections characteristic of OctaStar@Trp in PXRD spectra remain evident in the OctaStar@Trp+Alg material, indicating preservation of the underlying crystalline phase (Figure 2A). Such behavior is consistent with numerous biomolecule@MOF systems in which biomolecules are encapsulated, adsorbed, or immobilized within preformed frameworks while the underlying crystalline architecture remains intact.^48–49^ The incorporation of each component within the corresponding composite was examined using Fourier transformed Infrared (FTIR) (Figure 3A). The spectra of OctaStar, OctaStar@Trp, and OctaStar@Trp+Alg reveal that post-synthetic incorporation of Trp introduces only modest spectral changes while largely preserving the characteristic vibrational fingerprint of the parent OctaStar framework. The absence of major shifts, disappearance of framework-associated bands, or emergence of new coordination-related features indicates that Trp incorporation does not substantially alter the underlying Zn-imidazolate coordination network, consistent with guest confinement rather than framework reconstruction. Following alginate fixation, the principal vibrational features of the OctaStar framework remain largely unchanged (Figure 3A). At the same time, the appearance of a broad absorption centered at approximately 3300 cm^-1^, corresponding to O–H stretching vibrations from the abundant hydroxyl groups of the guluronic and mannuronic acid units of alginate,^50^ provides clear evidence for successful incorporation of the alginate into the composite. The coexistence of framework-associated vibrations together with the characteristic hydroxyl absorption supports the formation of the hierarchical OctaStar@Trp+Alg structure without disrupting the crystalline host. These spectra further support incorporation of Trp within the OctaStar host (Figure 3A). OctaStar exhibits a characteristic aliphatic C–H stretching vibration near 2884 cm^-1^ together with several fingerprint-region bands at approximately 1308, 1181, 1147, and 995 cm^-1^, which are characteristic of Zn-imidazolate frameworks and arise primarily from imidazole ring vibrations, including C–N stretching as well as C–C and C–H bending and rocking modes (Figure 3A).^51–52^ These framework-associated features remain clearly observable following Trp loading, further supporting preservation of the Zn-imidazolate network. In contrast, OctaStar@Trp exhibits an additional absorption near 2980 cm^-1^ assigned to C–H stretching vibrations associated with the tryptophan side chain, providing further evidence for successful incorporation of the Trp into the preformed framework (Figure 3A). Unlike the post-synthetic Route A materials (Figure 3A), the FTIR spectrum of the Route B Trp-MOF differs substantially from that of the parent Zn-imidazolate framework and exhibits features dominated by the coordination environment of Trp (Figure S2). The broad absorption centered near 3397 cm^-1^ is assigned primarily to hydrogen-bonded N–H and O–H stretching vibrations, while the band near 738 cm^-1^ corresponds to aromatic C–H out-of-plane bending associated with the indole ring of tryptophan (Figure S2). Together with the single-crystal X-ray diffraction (Figure S1), these spectral changes indicate that Trp participates directly in coordination during material formation, producing a distinct Zn–Trp coordination framework.

**Figure 3.**
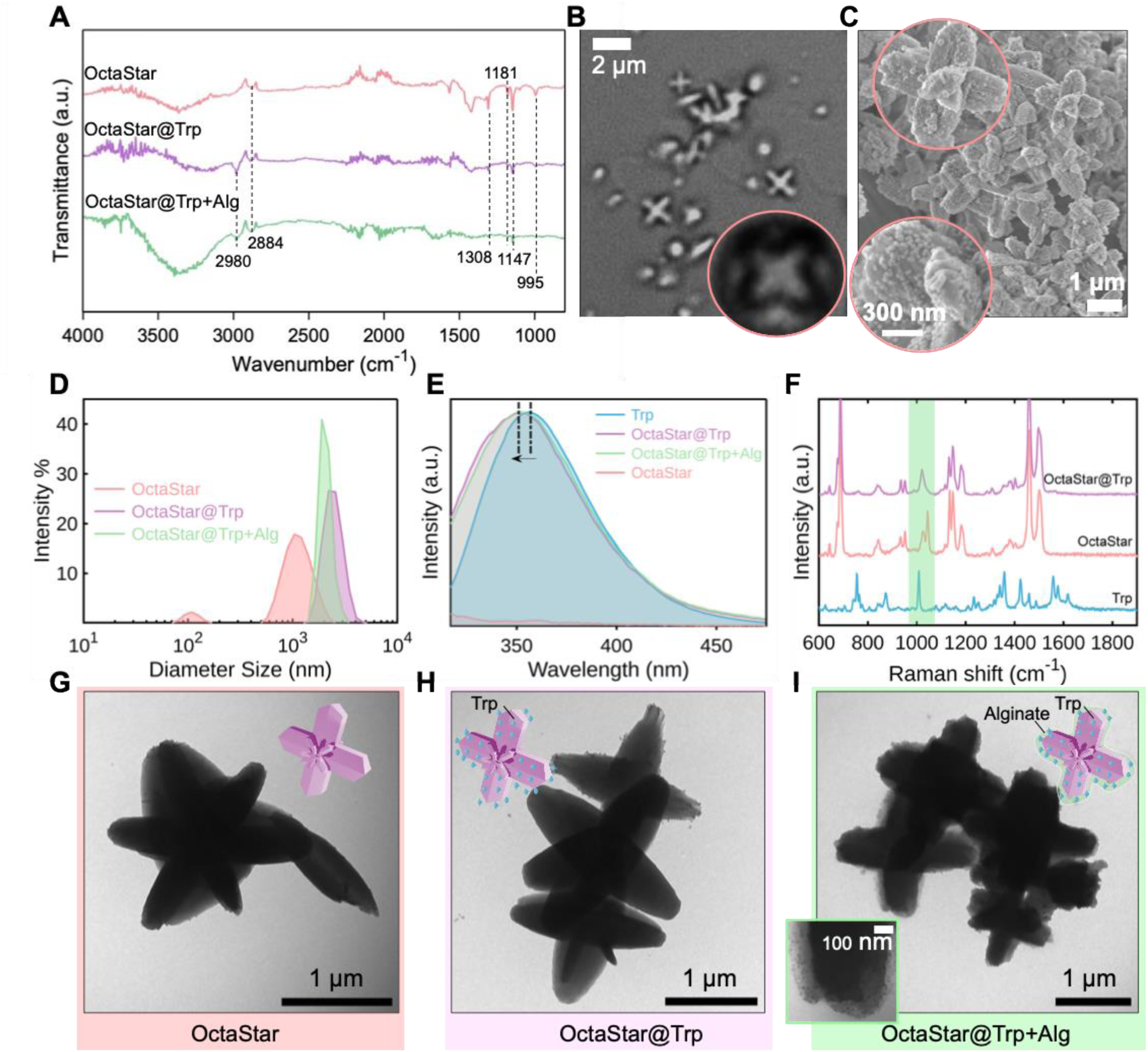
A) Fourier-Transformed Infrared (FTIR) spectra of the Trp, OctaStar, OctaStar@Trp, and OctaStar@Trp+Alg. B) Brightfield microscopy image of OctaStars. C) SEM image of OctaStar@Trp. D) Dynamic Light Scattering (DLS) size analysis of OctaStars, OctaStar@Trp, and OctaStar@Trp+Alg. E) Normalized Fluorescence Emission spectra of OctaStars, OctaStar@Trp, and OctaStar@Trp+Alg in comparison with free Trp (λ_ex_ = 306 nm). F) Raman spectra of the OctaStar in comparison with OctaStar@Trp and free Trp. G) TEM images of OctaStars, H) OctaStar@Trp, and I) OctaStar@trp+Alg.

Bright-field microscopy reveals the characteristic star-like morphology of the parent OctaStar particles (Figure 3B). SEM images demonstrate the highly faceted surface composed of interconnected plate-like crystallites that define the distinctive star-shaped architecture (Figure 3C). TEM images further corroborate these observations, showing that the characteristic multi-armed star morphology remains clearly identifiable after Trp loading (Figure 3G-H). Although alginate incorporation produces a slightly less well-defined particle outline (Figure 3I), likely reflecting deposition of the alginate within or around the OctaStar voids, the overall star-like architecture remains preserved. Dynamic light scattering (DLS) measurements indicate the main population of OctaStars around 1160 nm in size and remain within the same regime following each modification step (Figure 3D). The modest increase in apparent particle size following Trp incorporation and alginate treatment is consistent with formation of a guest-containing surface or interfacial layer while preserving the parent framework dimensions (Figure 3D). Unlike the star-like OctaStar particles observed in Route A, the SEM images of Route B material exhibit sheet-like and layered morphologies consistent with directional growth along preferred crystallographic planes (Figure S3). Such layered growth is highly consistent with the coordination geometry observed in the crystal structure (Figure S1), where Zn–Trp coordination propagates preferentially along 2D directions while extended aromatic interactions between indole groups further stabilize the layered architecture. Aromatic amino acids such as tryptophan are known to participate in strong π–π interactions and supramolecular ordering phenomena capable of influencing crystal packing and directional assembly behavior.^53^

Thermogravimetric analysis was used to compare the thermal behavior of the Route B Trp-MOF with that of ZIF-8 and free Trp (Figure S5). Free Trp exhibits a pronounced decomposition event beginning at approximately 280 °C, followed by rapid mass loss between approximately 300 and 430 °C and a residual mass of only ∼13% at 800 °C (Figure S5). In contrast, the Trp-MOF displays a broader, multistep mass-loss profile. Its principal decomposition begins near 280 °C, producing a gradual decrease to approximately 65% of the initial mass by ∼450 °C, followed by slower mass loss at higher temperatures. The Trp-MOF retains approximately 54% of its initial mass at 800 °C, a value comparable to the high-temperature residue observed for ZIF-8 and substantially greater than that of free Trp (Figure S5). The broader decomposition interval and increased residual mass indicate that incorporation of Trp into the extended Zn-coordination lattice alters its thermal decomposition pathway and provides greater thermal stabilization than is observed for the free amino acid. The distinct profile relative to ZIF-8 further supports the formation of a new Zn–Trp coordination phase rather than conventional ZIF-8 containing physically adsorbed Trp (Figure S5). Gas-sorption measurements were used to evaluate the accessible surface area and pore structure of the layered Route B Trp-MOF (Figure S6). Multipoint BET analysis over the selected linear relative-pressure range yielded a specific surface area of 70.5 m^2^ g^-1^ (Figure S6), demonstrating that the material possesses an appreciable adsorbate-accessible surface despite the relatively dense packing expected for an amino-acid-based coordination solid. The adsorption–desorption isotherm exhibits gradual uptake over most of the relative-pressure range, with only limited separation between the adsorption and desorption branches, followed by a sharp increase in uptake as (P/P_0_) approaches 1 (Figure S6). The high-pressure uptake is consistent with condensation within larger mesoporous or interparticle voids rather than uniform filling of a highly microporous three-dimensional framework. The absence of a pronounced low-pressure uptake further suggests that microporosity makes only a limited contribution to the measured surface area. The calculated pore-size distribution supports this interpretation, revealing accessible pores primarily between approximately 30 and 46 Å (Figure S6). These dimensions fall within the mesoporous range. Because single-crystal X-ray diffraction identifies Trp-MOF as a layered coordination structure rather than a conventional 3D porous ZIF topology, the measured mesoporosity may originate from accessible interlayer regions, structural defects, particle boundaries, or textural voids formed between the plate-like crystallites observed by SEM (Figure S3).

Because tryptophan is highly sensitive to its local molecular environment, fluorescence spectroscopy provides an additional probe of successful guest incorporation (Figure 3E). OctaStar does not show any fluorescence capability (Figure 3E), which is consistent with the fluorescence microscopy data (Figure 4). Whereas free Trp exhibits its characteristic emission maximum near 356 nm, encapsulated Trp within OctaStar displays a modest blue shift in emission to 351 nm (Figure 3E). Such behavior is consistent with confinement of Trp within a less polar and more spatially restricted microenvironment relative to bulk aqueous solution, where reduced solvent accessibility and restricted molecular motion can alter the excited-state relaxation pathway.^54^ Following alginate incorporation, the emission maximum remains largely preserved, indicating that the local environment experienced by the encapsulated Trp is maintained. These observations suggest that the OctaStar framework provides a stable host environment that governs the photophysical behavior of the confined amino acid while remaining compatible with subsequent hydrogel stabilization (Figure 3E). Raman spectra further confirm the presence of Trp within the OctaStar framework (Figure 3F). In addition to the characteristic vibrational features associated with the Zn-imidazolate network in OctaStars, OctaStar@Trp exhibits faint Trp features across the full 600-1800 cm^-1^ range and marked broadening of bands within the spectral region characteristic of indole ring vibrations, ∼950-1050 cm^-1^, providing further evidence for successful incorporation of Trp into the framework (Figure 3F). Importantly, the simultaneous observation of both OctaStar framework and Trp-associated Raman bands further supports guest confinement rather than replacement of the host structure, consistent with the FTIR and PXRD analyses.

**Figure 4.**
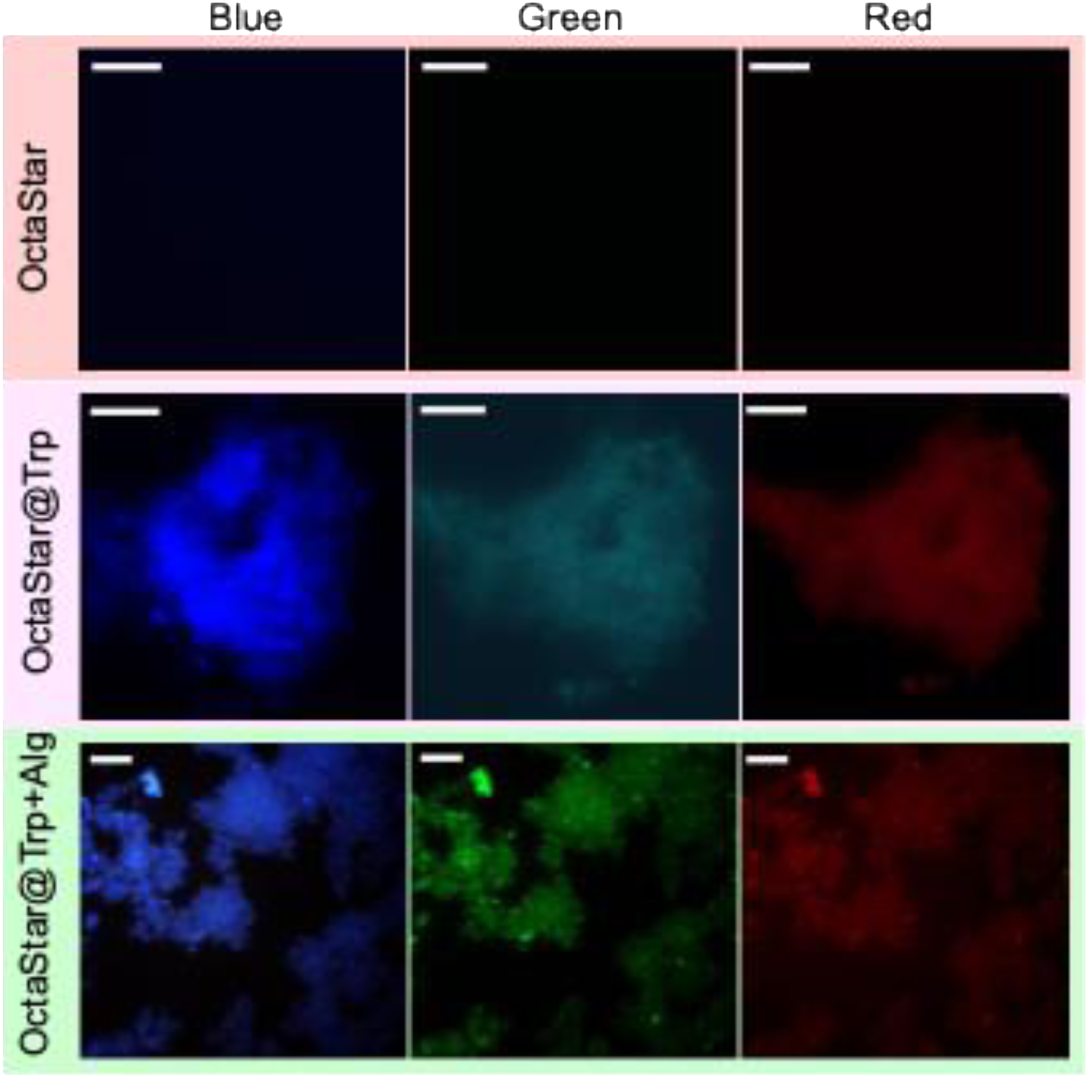
Fluorescence Microscopy images of OctaStar, OctaStar@Trp, and OctaStar@Trp+Alg., excited using blue (350 nm), green (488 nm), and red (561∼594 nm) light. Scale bars: 2 μm.

Fluorescence microscopy further verifies successful incorporation of Trp into the OctaStar framework (Figure 4). As expected, the parent OctaStar particles exhibit no fluorescence under all excitation channels examined, confirming that the Zn-imidazolate framework itself does not contribute significant intrinsic emission. In contrast, OctaStar@Trp displays strong fluorescence, providing direct visual evidence that Trp is successfully associated with the framework. The fluorescence is distributed throughout the particle rather than appearing exclusively at isolated surface locations, consistent with incorporation of Trp throughout the porous framework (Figure 4). Since Trp fluorescence is highly sensitive to local polarity, solvent exposure, electrostatic environment, and conformational dynamics,^17, 19^ the photophysical response of OctaStar@Trp could be due to the changes in local polarity, hydration, molecular mobility, and diffusional exchange. Following alginate fixation, the fluorescence remains clearly associated with the particles, demonstrating that the alginate fixation process does not remove or substantially quench the incorporated Trp. Instead, the OctaStar@Trp+Alg particles continue to exhibit strong fluorescence (Figure 4). These results therefore support a hierarchical host–guest structure in which Trp remains confined within a structurally preserved OctaStar framework while the alginate network provides further stabilization without compromising the optical properties of the encapsulated amino acid. Interestingly, Trp-MOF synthesized via Route B was also fluorescent under DAPI, GFP and TXRed showing that the coordination of tryptophan to the zinc centers induce fluorescence across the visible range (Figure S4). Since Trp is directly integrated into the coordination lattice in Route B, the local environment surrounding the indole fluorophores becomes substantially more ordered and rigid relative to the guest-confined Route A materials. The layered structure likely introduces extensive intermolecular interactions between neighboring Trp residues together with reduced solvent accessibility, both of which are known to strongly influence indole fluorescence behavior.^54^ It has been previously shown that the optical properties of organic molecules in crystalline form are intimately linked to their crystal macrostructure and the relative spatial arrangement of those molecules across many length scales within the crystal.^55^ Three kinds of inter- or intrachain interactions, including hydrogen bonding, π–π and n−π coupling, have been suggested to contribute to fluorescence emission of materials in the rigid conformation.^54^ Thus, incorporation of fluorescent guests into MOFs is a particularly active area due to their ordered porous environment that can rigidify the guest, suppress aggregation-related quenching, and modulate emission through host–guest interactions.^54, 56–58^ Consequently, MOFs are often treated as structurally passive hosts into which photophysically active molecules are incorporated after framework formation. Biomolecules, however, are not always well described as inert guests. Amino acids, peptides, and many cofactors carry carboxylate, amine, imidazole, indole, or hydroxyl groups that can coordinate Zn^2+^ and related metals.^25, 30, 59^ In this work, the fluorescence of OctaStar@Trp is highly suitable for biological and medical applications due to their biocompatibility and ease of preparation.^60–61^

Bright-field microscopy demonstrates that the characteristic star-shaped morphology of the OctaStar particles remains readily identifiable following both Trp incorporation and subsequent alginate fixation (Figure 5A, B, & C). These observations agree well with the SEM and TEM analyses presented previously, indicating that the hierarchical assembly process preserves the overall architecture of the OctaStar host (Figure 3). Post-synthetic loading of Trp into the OctaStar framework yields an encapsulation efficiency of approximately 51%, confirming that the porous OctaStar host readily accommodates the amino acid (Figure 5D). Following alginate incorporation, the measured encapsulation efficiency decreases modestly to ∼37% (Figure 5D). This reduction is potentially associated with the additional processing required for alginate gelation and the formation of the polymer network, during which a fraction of loosely associated Trp may be lost prior to complete Ca^2+^-induced crosslinking. Importantly, despite this modest decrease in loading, a substantial amount of Trp remains associated with the composite, demonstrating that the alginate immobilization process is compatible with efficient guest incorporation while simultaneously introducing an additional level of structural stabilization. Direct incorporation of Trp throughout the coordination lattice in Route B produces ∼85% Trp encapsulation efficiency, which is substantially higher than post-synthetic diffusion-based guest uptake into a pre-existing framework. The release profiles reveal the complementary roles of guest confinement and secondary alginate fixation (Figure 4F-G). OctaStar@Trp exhibits rapid release under acidic conditions, with nearly three-quarters of the encapsulated Trp released within the first hour at pH 3, whereas considerably lower release is observed at neutral and basic pH (Figure 5E). This behavior is consistent with the well-established acid sensitivity of Zn-imidazolate frameworks. With 2-methylimidazole as the organic linker (pKa = 7.87),^62^ the framework relies on the deprotonated 2-methylimidazolate anion to maintain coordination with Zn centers. Under acidic conditions, protonation of the linker weakens the Zn–imidazolate coordination network, promoting framework degradation and facilitating diffusion of the confined Trp into the surrounding solution.^59, 63–66^ In contrast, OctaStar@Trp+Alg exhibits substantially lower cumulative release under all investigated pH conditions. Even under strongly acidic conditions, where partial degradation of the Zn-imidazolate framework is expected to occur, Trp release remains markedly suppressed relative to the non-gelled material, while only minimal release is observed under neutral and basic conditions. These results demonstrate that alginate fixation effectively enhances guest retention without eliminating the environmentally responsive nature of the release process. Unlike the Zn-imidazolate framework, the Ca^2+^-crosslinked alginate network remains structurally intact under mildly acidic conditions owing to its stable cooperative coordination between calcium ions and the guluronate-rich blocks of the polysaccharide.^67^ Rather than undergoing complete dissolution, calcium alginate hydrogels have been shown to shrink and deform upon acidification,^68–70^ forming a denser polymeric network that can further restrict molecular diffusion. Such behavior is consistent with our proposed localized gelation mechanism, where the alginate network forms a secondary interconnected matrix within and around the OctaStar voids, thereby introducing an additional physical barrier that limits Trp diffusion even as the underlying Zn-imidazolate framework begins to degrade (Figure 5E). This hierarchical structure therefore establishes two sequential barriers to mass transport. The OctaStar framework serves as the primary host responsible for molecular confinement of Trp, while the crosslinked alginate network functions as a secondary diffusional barrier that continues to regulate guest transport after partial framework degradation. Consequently, release is no longer governed solely by the stability of the Zn-imidazolate framework but also by diffusion through the surrounding alginate matrix. This dual-barrier mechanism explains the substantially improved retention observed for OctaStar@Trp+Alg across all investigated pH values. The experimental release profiles further support this mechanism (Figure 5E). Both composites exhibit an initial burst release during the first 15 min at pH 3 and 5, followed by a gradual approach to equilibrium, whereas considerably slower release is observed over the three-hour period at pH 9, consistent with the greater stability of Zn-imidazolate frameworks under basic conditions (Figure 5E). The release curves subsequently plateau, indicating that little additional Trp is liberated after the initial diffusion-controlled stage. Importantly, incorporation of alginate dramatically reduces the total amount of Trp released at every pH investigated (Figure 5E). The cumulative release decreases from ∼ 78%, 75%, and 64% for OctaStar@Trp to ∼ 38%, 33%, and 22% for OctaStar@Trp+Alg at pH 3, 5, and 9, respectively (Figure 5E). While alginate reduces Trp release by approximately 2-fold under acidic conditions, the reduction approaches 3-fold under basic conditions, reflecting the combined effects of an intact Zn-imidazolate host and the additional alginate diffusion barrier (Figure 5F-G).

**Figure 5.**
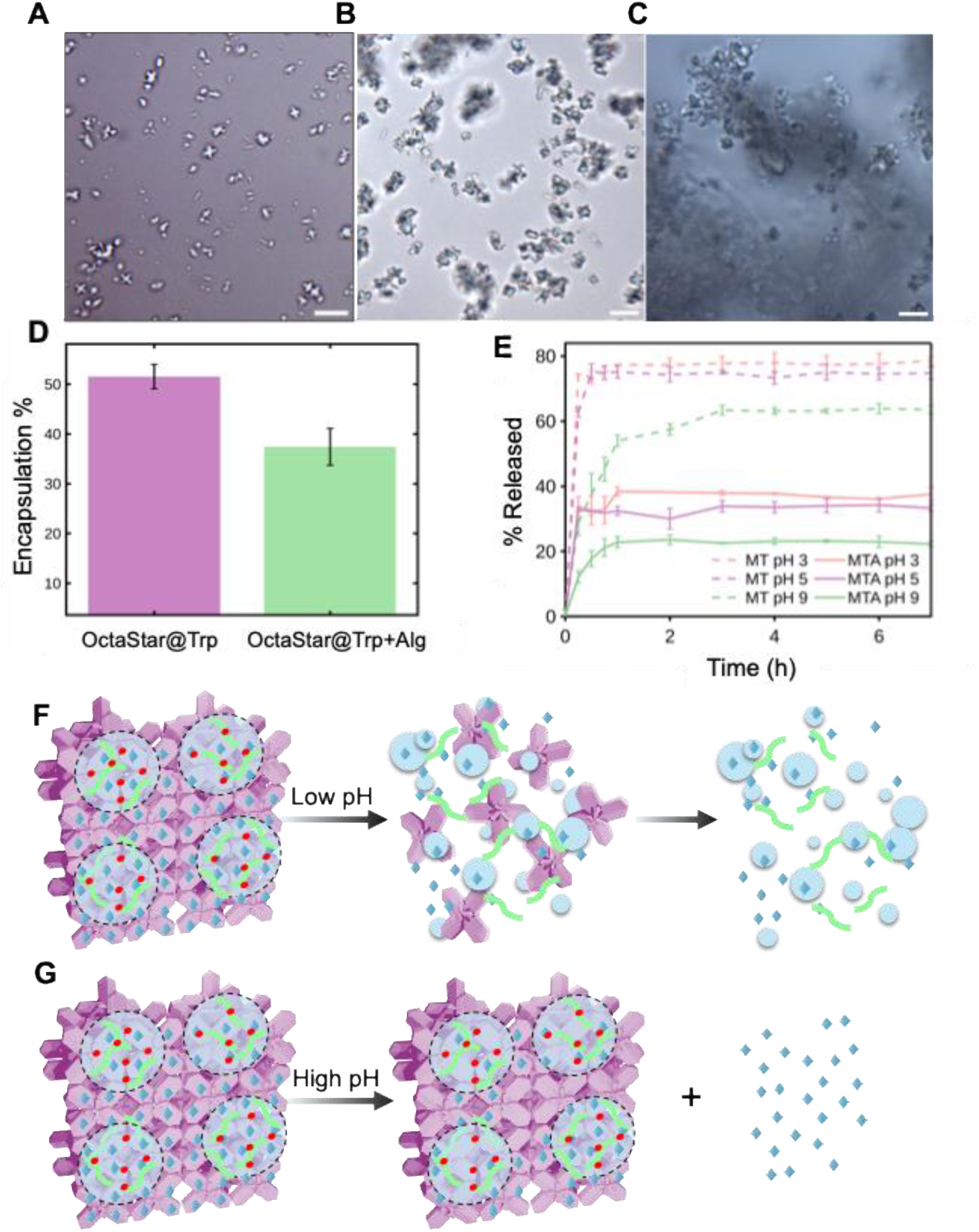
Brightfield microscopy images of A) OctaStars, B) OctaStar@Trp, and C) OctaStar@Trp+Alg. D) Encapsulation efficiency of tryptophan in OctaStar@Trp in comparison with OctaStar@Trp+Alg. E) The release percentage of Trp from OctaStar@Trp and OctaStar@Trp+Alg at pH 3, 5, and 9. F) The schematic of OctaStar@Trp+Alg dissociation in low pH and G) high pH environment.

The comparison between these two assembly pathways reveals that Trp cannot be regarded simply as molecular cargo introduced into an otherwise passive host. Its function within the material depends on the structural role it assumes during crystallization and on the molecular environment created by that role. When confined as a guest, its accessibility remains governed by the stability and transport properties of the surrounding host, whereas coordination-driven integration couples its behavior directly to the organization of the crystalline lattice. Thus, the relationship between assembly pathway, structural role, and local environment provides the unifying framework through which the distinct behaviors of these materials can be understood.

## CONCLUSION

In this work, we demonstrate that the structural organization of tryptophan within crystalline materials can be deliberately controlled through selection of the assembly pathway. Although both routes employ Zn^2+^, 2-methylimidazole, and Trp, introducing the amino acid before or after framework formation produces two fundamentally different materials. Post-synthetic loading confines Trp as a guest within a preformed star-shaped Zn–imidazolate host while largely preserving the parent crystalline architecture. In contrast, simultaneous combination of the components causes Trp to participate directly in coordination-driven assembly, producing a distinct layered Zn–Trp crystalline phase in which 2-methylimidazole appears to act as a transient mediator rather than a permanent framework linker. The resulting organization modes provide complementary functional characteristics. Guest confinement in the OctaStar host preserves the intrinsic fluorescence of Trp and produces pH-responsive release governed by the environmental stability of the Zn–imidazolate framework. Secondary calcium–alginate fixation introduces an additional diffusional barrier that substantially improves Trp retention without disrupting the crystalline host or eliminating its fluorescence. Direct coordination-driven integration in Route B provides higher Trp incorporation, a thermally stabilized coordination lattice, accessible solid–gas interfaces, and a strong optical response under multiple excitation conditions. Together, the crystallographic, spectroscopic, microscopic, thermal, adsorption, and release results show that these materials cannot be described simply as different carriers containing the same molecular cargo. Rather, their properties emerge from the specific structural role assigned to Trp during assembly. These findings establish synthetic pathway selection as a powerful means of controlling whether a biologically important amino acid behaves as a confined guest or as an integral structural component of a crystalline material. More broadly, the study illustrates how organization across molecular and crystalline length scales can regulate optical behavior, molecular accessibility, thermal stability, and guest retention without changing the underlying molecular building blocks. This organization-centered strategy provides a foundation for designing amino-acid-based materials for responsive fluorescence sensing, controlled molecular delivery, and biomimetic systems in which the spatial arrangement of redox-active residues is engineered to support directional charge or energy transfer.

## Acknowledgements

We acknowledge support from the College of Arts & Sciences Microscopy and X-ray Diffraction Facility. We are grateful for the funding support from the ACS Petroleum Research Fund Award 69034-DNI3.

## Data Availability Statement

The supplementary information, raw data and custom analysis scripts are available upon request from the corresponding author.

