## Supporting Information PDF for "Engineering the Structural Organization of Tryptophan in Crystalline Materials for Tunable Functionality"

### MATERIALS AND METHODS

Materials

L-Tryptophan was purchased from Thermo Fischer Scientific (Waltham, MA, USA). Zinc nitrate hexahydrate, 2-methylimidazole, and alginic acid sodium salt were purchased from Sigma Aldrich (St. Louis, MO, USA). Calcium chloride dihydrate was purchased from Fisher Scientific (Hampton, NH, USA). DI water was filtered using PURELAB Flex 2 system from Evoqua Water Technologies (Pittsburgh, PA, USA)

Experimental Methods

Synthesis of OctaStars

The first stock solution was prepared by dissolving 1.86 g zinc nitrate hexahydrate in 50mL DI water to form a 0.125M solution. The second solution was prepared by dissolving 8.21 g of 2-methylimidazole in 100 mL of DI water to make 1M solution. Then, 2 mL aliquot of MeIm solution was added to a 3.5 mL glass vial, continued 1mL aliquot of zinc solution under stirring at 1500 rpm. The solution immediately became cloudy and was stirred for 30 minutes. Once taken off the stir plate, it sat for another 30 minutes before being centrifuged at 14,000 rpm for 30 minutes. The pellet was then washed with 3 mL DI water to separate the residual precursors.

Synthesis of OctaStar@Trp via incorporation of Tryptophan

OctaStar pellet was resuspended using 1.5 mL Trp solution (10 mg/mL) and transferred to a falcon tube, to which 4.5 mL DI water was added. The solution was then stirred at 1500 rpm for 1 hour and centrifuged for 30 mins at 7,000 rpm. The final pellet was then washed with 3 mL of DI water.

Synthesis of OctaStar@ Trp+Alg via incorporation of Trp and sodium alginate

OctaStar pellet was resuspended using 1.5mL Trp solution (10mg/mL) and transferred to a falcon tube, to which 4.5mL sodium alginate solution (1% wt) was added. The solution was then stirred at 1,500 rpm for 1 hour and centrifuged for 30 mins at 7,000 rpm. The supernatant was removed and 3mL of Olive Oil was added. After stirring for 10 mins at 1,500 rpm, 3 mL of $CaCl_2$ was added slowly dropwise. After another 20 mins of constant stirring, the final solution was centrifuged at 7,000 rpm for 30 mins. To get solid particles out of the solution for characterization, pentane was added and the solution was stirred at 1,500 rpm for 10 minutes. Then, 3 mL of 2M $CaCl_2$ solution was added dropwise. After 5 minutes, the solution was slowly heated to 40 °C for 15 minutes to evaporate any remaining Pentane. Then, the solution was transferred to small Eppendorf tubes and centrifuged at 14,000 rpm for 30 minutes. The final pellet was then washed with 3 mL of DI water.

Synthesis of Trp-MOF

In a test tube, 2 mL of 50 mM L-Trp solution, 2 mL of 50 mM zinc nitrate solution and 2mL of 50 mM MeIM solution were mixed and stirred in a water bath at 50°C for 2 hours. The mixture was then centrifuged at 20,000 rcf, the supernatant was removed, the pellet was washed 3 times and dried using a Speed Vac.

Synthesis of ZIF-8

Stock solutions of 835 mM 2-methylimidazole and 101 mM zinc nitrate hexahydrate were prepared. Then, in a glass vial, 1.25mL DI water, 1.75 mL 2-methylimidazole, and 0.25mL of zinc nitrate were mixed respectively. After 3 hours, the mixture was centrifuged at 25,000 rcf for 20 minutes. The prepared ZIF-8 was washed with DI water and dried using Savant Speed Vac Plus SC110A from Savant Instruments Inc. Farmingdale, New York, USA.

Release Study

The obtained particles were resuspended and stirred at 1,000 rpm for 7 hours in 11 mL solution of 0.1 M buffers at different pHs (citrate buffer pH 3, acetate buffer pH 5, and Tris-HCl buffer pH 9. At each data point, 1 mL sample was aliquoted and centrifuged for 30 minutes at 14,000 rpm. The supernatant was collected and examined under UV-vis to monitor the release tryptophan.

Ultraviolet–visible (UV-vis) Spectrophotometry

UV-vis spectrophotometry was conducted in the range of 190-900 nm, with 1.0 nm data interval, and 0.004 s averaging time on an Agilent Cary UV-Vis Compact Peltier from Agilent Technologies, Mulgrave, Australia. The samples were in aqueous solution and measured using standard quartz cuvette.

Dynamic Light Scattering (DLS)

The size of synthesized particles was investigated by dynamic light scattering (DLS) mode of Malvern Zetasizer pro from Malvern Panalytical Ltd., Grovewood Road, Malvern, Worcestershire, WR14 1XZ, United Kingdom. This study was performed using disposable plastic cuvettes provided by Malvern. A fluorescence optical filter was used to improve detector's sensitivity.

Fluorescence Spectroscopy Measurements

Fluorescence Emission was obtained using Agilent Technologies Cary Eclipse Fluorescence Spectrometer at room temperature using standard quartz cuvettes with a 1 cm path length. All samples were excited at 306 nm using excitation and emission spectral band passes of 5 nm. The PMT voltage was set to "Medium" with a scan speed of 20 nm/s. The samples were in solution.

Attenuated total reflection Fourier transformed infrared spectroscopy (ATR-FTIR)

ATR-FTIR was performed on a Nicolet iS10 FT-IR with a Smart iTX ATR sampling accessory from ThermoFisher Scientific, Thermo Electron Scientific Instruments LLC, Madison, WI, USA. The samples were dried using Savant Speed Vac Plus SC110A from Savant Instruments Inc. Farmingdale, New York, USA.

Single-crystal X-ray diffraction

The crystal structure and unit cell were identified using a Rigaku Synergy-S equipped with both CuKα and MoKα radiation and paired with a Rigaku HyPix-6000HE detector. The phase composition and crystal structure were determined using a Rigaku Miniflex II which uses a CuKα radiation in a Bragg-Brentano geometry over a range of 3.0°-60°, with a scanning rate of 1°/minute. To grow Trp-MOF crystals, 6 mL of 50 mM L-Trp, 6mL of 50 mM zinc nitrate and 6 mL of 50 mM MeIM were mixed and then placed in a 50°C oven for 3 days. The crystals were filtered, washed and dried in a 50°C oven.

Powder X-ray diffraction (PXRD)

Crystalline structure of the synthesized MOFs was investigated using PXRD. Samples were dried using Speed Vac. The samples were then placed on a sample disk and PXRD patterns were recorded at a 5°/min scanning speed and 5−60° diffraction angle by a Rigaku MiniFlex II Powder X-Ray Diffractometer with Cu Ka radiation set at 30 kV and 15 mA from Rigaku Americas Corporation, 9009 New Trails Drive, The Woodlands, TX, USA. A monochromator filter was also used to reduce interferences from fluorescence.

Fluorescence Microscopy

Fluorescence microscopy images of Route A were taken using an Olympus BX41 clinical microscope, trinocular brightfield. The 4 light filter settings and their respective excitation wavelength were: 4',6-Diamidino-2-phenylindole - DAPI (350 nm), Green Fluorescent Protein - GFP (488 nm), Texas Red (561-594 nm). Fluorescence microscopy images of Route B were taken using a Leica THUNDER Imager Cell Spinning Disk with a widefield modality. The dried Trp-MOF sample was mounted to a glass microscope slide. The filters used were DAPI, GPF, TXRed, and Cyanine5 (Cy5, 647nm).

Brunauer–Emmett–Teller (BET) Analysis

Surface area analysis was performed using $N_2$ adsorption-desorption isotherms at 77K with a Quantachrome® ASiQwin™ analyzer. Samples were dried on a watch glass in a 50 °C oven for 3 days. Prior to measurements, samples were degassed under vacuum at 100°C for 12h to eliminate adsorbed gases and moisture. The specific surface area of the samples was determined using the Brunauer–Emmett–Teller (BET) method. The BET equation was applied in the relative pressure range ($P/P_0$) of 0.05–0.50, where the isotherms showed linearity. The surface area was calculated based on the measured adsorption data. Pore size distribution and pore volume were further analyzed using Density Functional Theory (DFT) applied to the N2 adsorption-desorption isotherms. The analysis was carried out using N2 at 77 K on carbon (cylindrical pores, QSDFT adsorption branch).

Scanning Electron Microscopy (SEM) and Transmission Electron Microscopy (TEM)

Surface morphology was investigated by scanning Electron Microscopy (SEM) using a Zeiss crossbeam 540 FIB-SEM along with the Hitachi S-4300 high resolution field emission SEM1. The SEM images were collected at 5 kV accelerating voltage and 5.9 mm working distance with side mount secondary electron detector. Samples were dropped onto aluminum sample stubs and were observed after drying. TEM was performed using a Hitachi S-7650 TEM.

Thermal Gravimetric Analysis (TGA)

Thermal stability of the samples was investigated using TGA. The samples (10 mg) were placed on a sample pan and heated in a Nitrogen atmosphere from 25 °C to 800 °C at a rate of 10 °C/min using a Shimadzu DTG-60H simultaneous DTA-TG apparatus from Shimadzu Corporation, Kyoto, Japan.

Confocal Raman Microscopy

Raman spectra were recorded using a confocal Raman microscope described in detail previously.[1-2] Samples were excited at 660 nm using 1 mW laser power measured at the back of the microscope. Samples were dispersed on the surface of a glass microscope coverslip (No. 1.5 thickness Corning BK7 glass, Fisher Scientific, Waltham, MA USA). Excitation radiation was focused within the sample from below using an oil immersion microscope objective (100 ×, Olympus, Waltham, MA, USA). The acquisition time for each spectrum was 60 s. Spectra were processed using custom MATLAB (R2022a; MathWorks, Natick, MA) scripts.

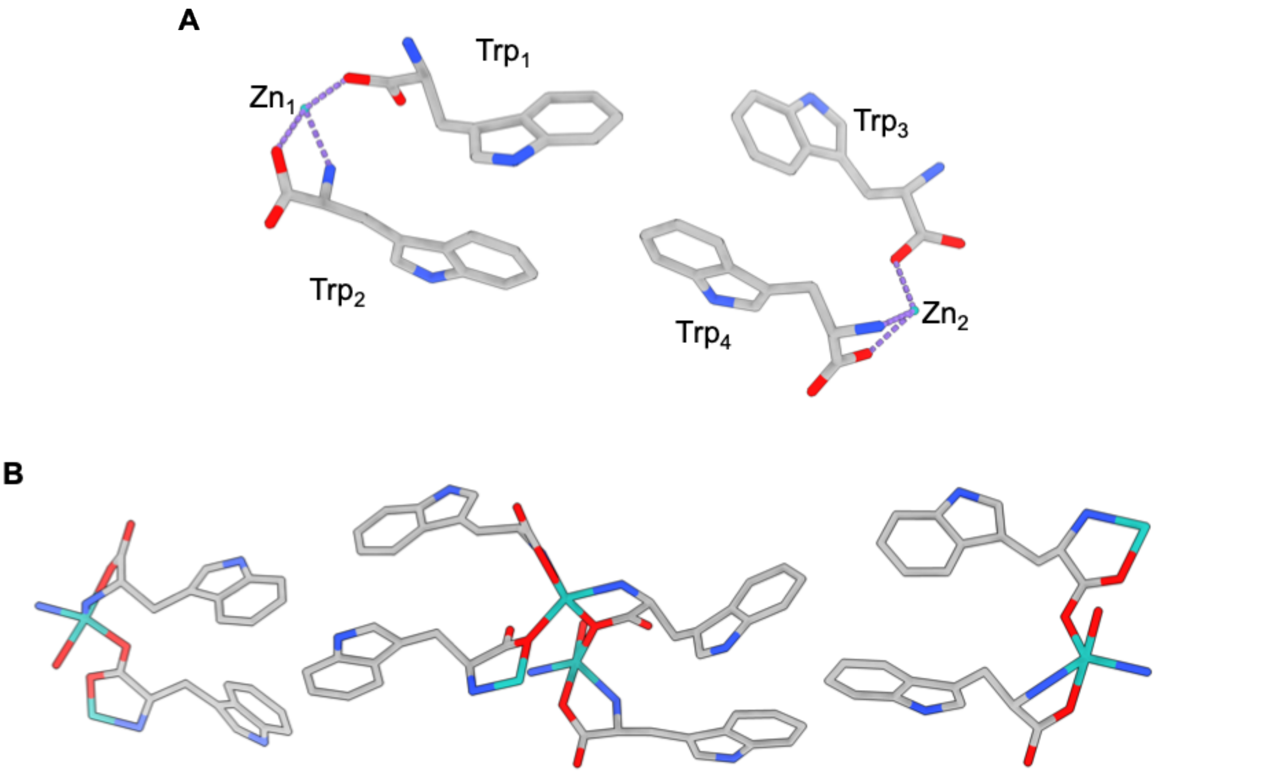


Figure S1. Molecular structure derived from single-crystal X-ray diffraction analysis of MOF synthesized through Route B. A) Simplified representation of the local coordination environment illustrating the coordination of Zn by two independent tryptophan ligands. The indole nitrogen atoms remain non-coordinating, with Zn binding occurring exclusively through the α-amino nitrogen and carboxylate oxygen atoms. B) Crystallographic coordination motifs highlighting the local Zn–Trp environments within the layered crystal architecture. Notably, although 2-methylimidazole was required during synthesis, it was not observed as a coordinated component in the final crystal structure, indicating its role as a structure-directing agent during assembly rather than a permanent framework ligand. Zn, cyan; O, red; N, blue; C, gray. Hydrogen atoms are omitted for clarity.

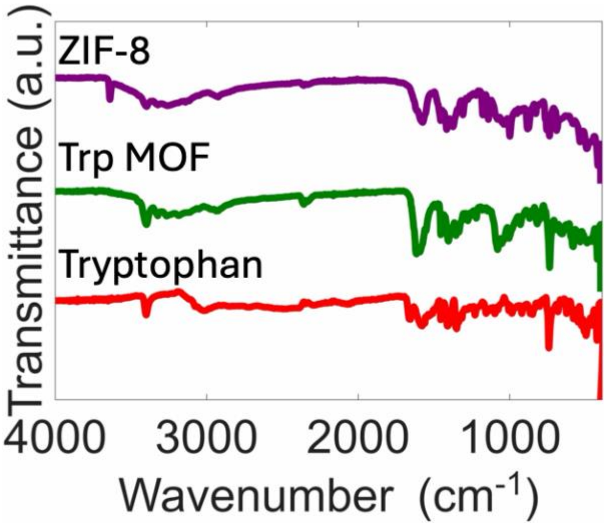


Figure S2. FTIR spectra of the Trp-MOF synthesize via Route B, ZIF-8, and free Trp.

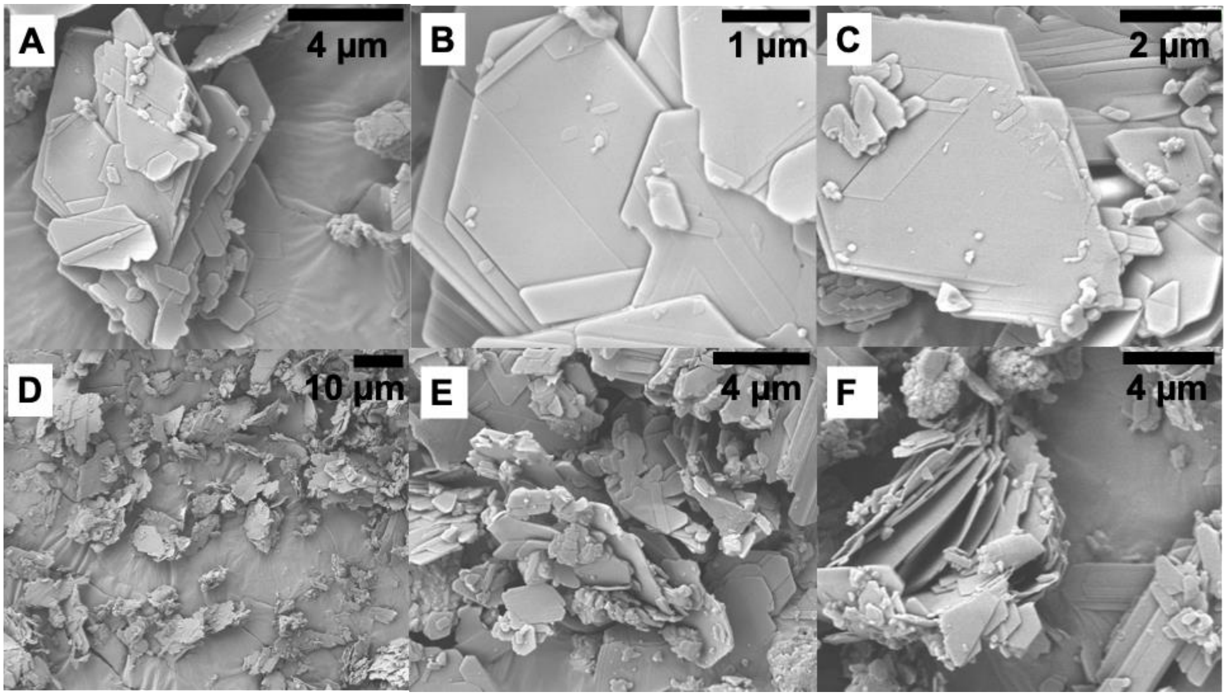


Figure S3. SEM images of Trp-MOF synthesized via Route B.

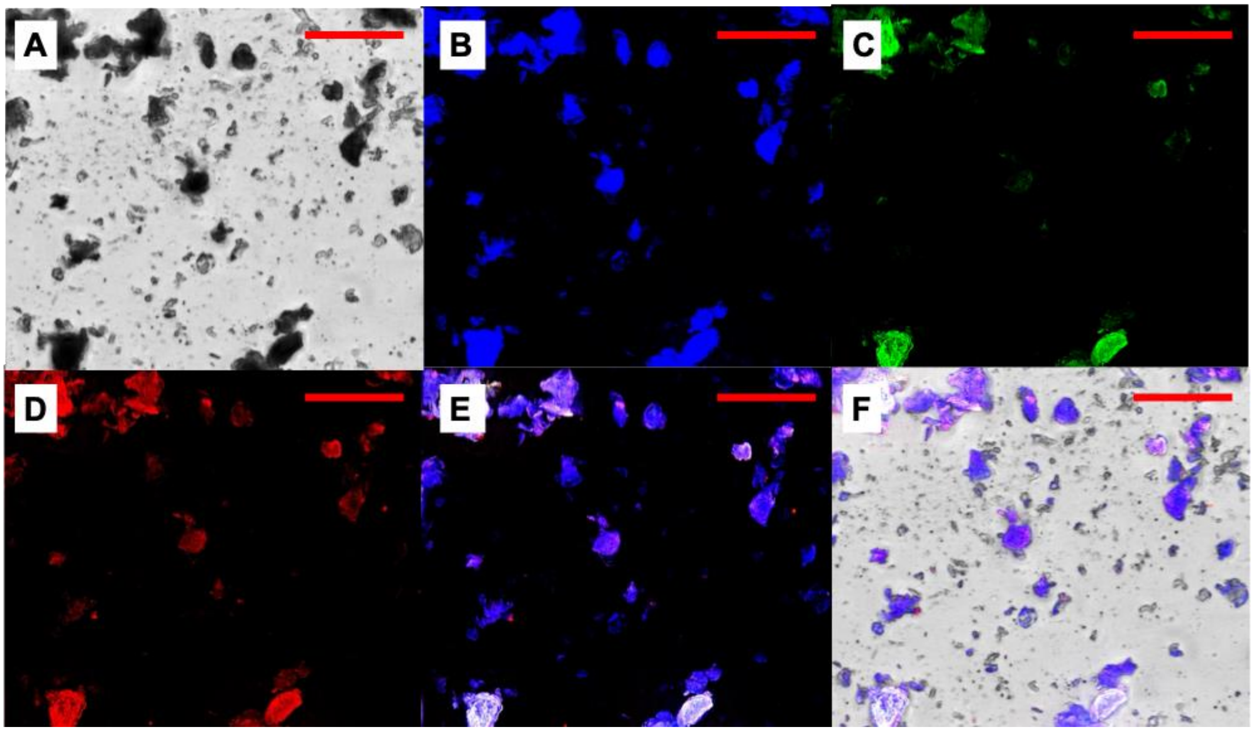


Figure S4. Fluorescence Microscopy images of Trp-MOF synthesized via Route B. A) Bright field, B) DAPI, C) GFP, D) TXRed Filters, E) overlay of DAPI, GFP and TXRed, and F) overlay of bright field, DAPI, GFP and TXRed. Scale bars: 30µm.

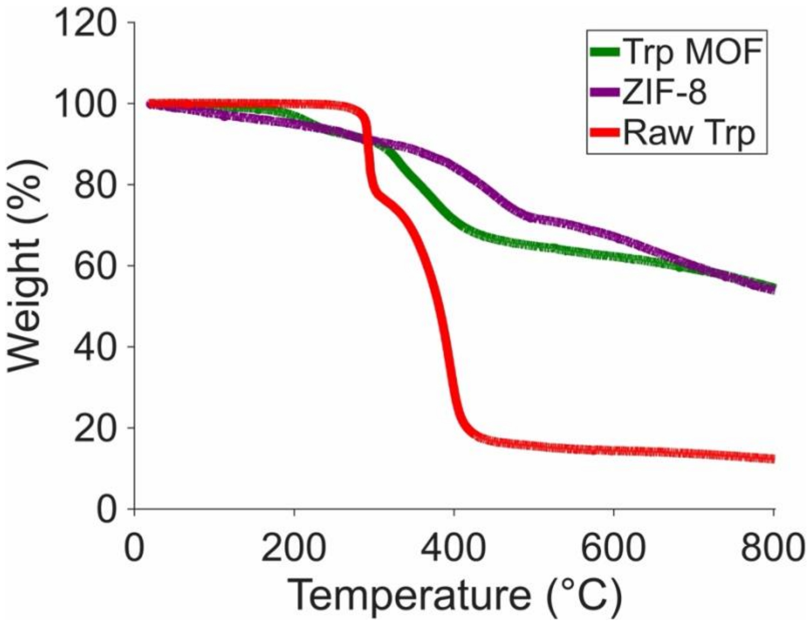


Figure S5. Thermogravimetric analysis (TGA) of Trp-MOF compared with ZIF-8 and free Trp.

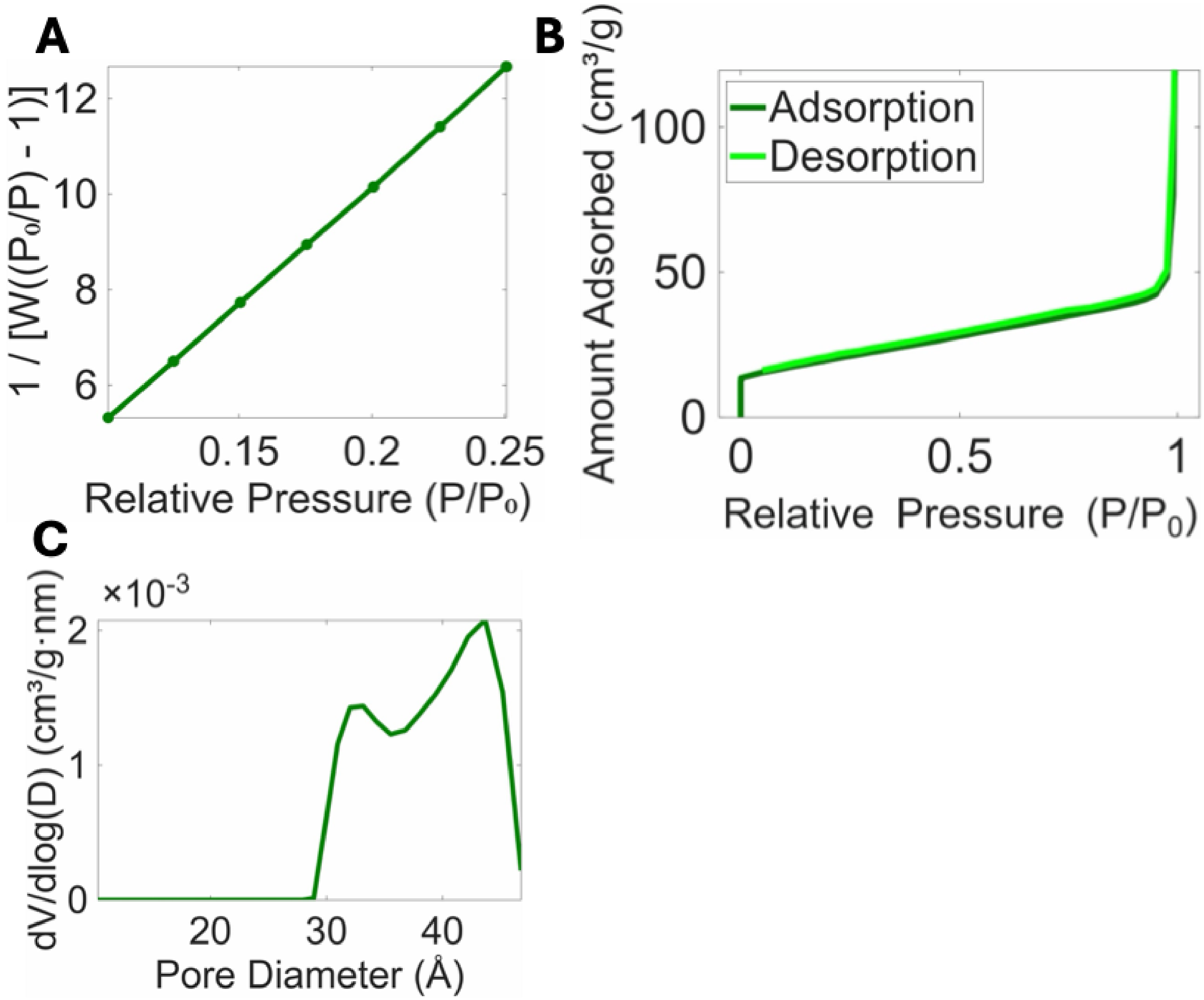


Figure S6. Surface-area and pore-structure characterization of the Route B Trp-MOF. A) Multipoint Brunauer–Emmett–Teller (BET) plot over the selected relative-pressure range used to calculate a specific surface area of 70.5 $m^2$ $g^{-1}$. B) Gas adsorption–desorption isotherms of Trp-MOF plotted as the amount adsorbed as a function of relative pressure, ($P/P_0$). C) Pore-size distribution calculated from the desorption branch of the isotherm, showing pores primarily within approximately 30–46 Å, consistent with mesoporous domains in the material.

Table 1. Single-crystal X-ray diffraction analysis for Trp-MOF formed via Route B

| | |
|---|---|
| Crystal System | Monoclinic |
| Space Group | $P2_1$ |
| a (Å) | 9.3060 |
| b (Å) | 5.4349 |
| c (Å) | 12.9767 |
| α (°) | 90° |
| β (°) | 91.548° |
| γ (°) | 90° |

1. Xu, J.; Koh, M.; Minteer, S. D.; Korzeniewski, C., In situ confocal Raman microscopy of redox polymer films on bulk electrode supports. *ACS Measurement Science Au* **2023,** *3* (2), 127–133.
2. Pasikku Hannadige, L.; Kusoglu, A.; Korzeniewski, C., Confocal Raman Microscopy as an In Situ Probe of Volume Change in Hydration-Sensitive Polymer Membranes. *Analytical Chemistry* **2025,** *97* (27), 14126–14131.
